# MtbAIM: AI/ML-guided endpoint-specific prediction of small-molecule activity across drug-susceptible and drug-resistant *Mycobacterium tuberculosis*

**DOI:** 10.64898/2026.09.28.754881

**Authors:** Jasdeep Singh, Kritika Singh

**Author notes:** Correspondence: Jasdeep Singh.

## Abstract

The emergence of drug resistance in *Mycobacterium tuberculosis* (M.tb) presents a major challenge to drug discovery, as compounds active against drug-susceptible (DS) isolates may not retain comparable inhibitory activity against drug-resistant (DR) isolates. Here, we developed an AI/ML workflow to independently prioritize small molecules separately against DS and DR isolates of M.tb and predict MIC, MIC90, and MIC99 as distinct quantitative endpoints. Following curation of data retrieved from the ChEMBL repository, molecular-activity datasets comprised 25,645 and 3,749 compounds tested against whole cell DS and DR isolates respectively. Molecular representations were generated using RDKit descriptors and Morgan fingerprints, along with embeddings from ChemBERTa, MolCLR, and Uni-Mol, and evaluated for binary activity classification and endpoint-specific regression. The best-performing DS classifier combined RDKit/Morgan features with ExtraTrees (balanced accuracy 0.782; AUROC 0.864; AUPRC 0.870), while histogram gradient boosting using the same features performed best for DR classification (balanced accuracy 0.762; AUROC 0.836; AUPRC 0.880). Similarly, RDKit/Morgan features combined with tree-based regressors achieved the best overall performance across all six DS/DR regression endpoints, with pooled out-of-fold RMSE values ranging from 0.604 to 0.784 pMIC units. To facilitate broader application, we developed MtbAIM (https://github.com/jasdeep002/MtbAIM), a Google Colab–based prediction platform that enables users to assess and prioritize small molecules against DS and DR isolates of M.tb.

## Introduction

Tuberculosis remains one of the most persistent infectious diseases globally. The World Health Organization estimated that 10.7 million people developed tuberculosis in 2024 and approximately 1.23 million died from the disease in the same^1^. Drug resistance makes this burden harder to control by narrowing treatment options and also increasing economic burden, need for compounds with new chemistry and mechanisms of action^2^. For drug discovery, a consistent challenge is not only whether a molecule could M.tb, but also retention of activity against resistant strains. Classical drug discovery approaches may cost millions, even before a drug reaches clinical trials.

AI and machine learning (ML) have now expanded our potential to explore vast chemical space before expensive experimental testing. The discovery of anti-mycobacterial compound Halicin provided an early example of how learned molecular representations can identify candidates outside conventional screening priorities^3^. In tuberculosis research, ML models have since been developed from curated antitubercular datasets, ChEMBL-derived activity records, fingerprints, molecular descriptors, and ensemble classifiers^4-6^. These studies established that chemical structure carries useful information for antitubercular prediction, however a critical gap remains–majority of these workflows emphasize targeting wild-type H37Rv, pooled MIC measurements, or a single active/inactive outcome^7-9^.

To address this gap, we designed an analytical framework to prioritize small molecule(s) with anti-mycobacterial potential, separately against drug-susceptible and drug-resistant isolates while preserving the identity of individual activity endpoints. To achieve this, we modeled DR and DS records separately, and MIC, MIC90, and MIC99 were retained as independent classification and regression targets. We then compared conventional cheminformatics features (RDKit, Morgan fingerprints) along with three learned molecular representations (ChemBERTa^10^, MolCLR^11^ and UniMol^12^) under the same scaffold-aware validation framework. This design allowed us to ask two related questions: how well anti-tubercular activity generalizes to unseen chemical scaffolds, and whether modern learned representations improve that generalization over established RDKit/Morgan features^13^. An overview of the workflow used in this study is outlined in Figure 1. To the best of our knowledge, this study represents one of the first pilot-scale AI/ML frameworks for small-molecule antitubercular discovery to explicitly stratify DS and DR M.tb data while performing endpoint-specific binary classification and regression. The final prediction platform, developed as MtbAIM is freely accessible through Google Colab (https://github.com/jasdeep002/MtbAIM), enabling users to perform predictions without requiring local software installation or committing huge computational resources.

**Figure 1.**
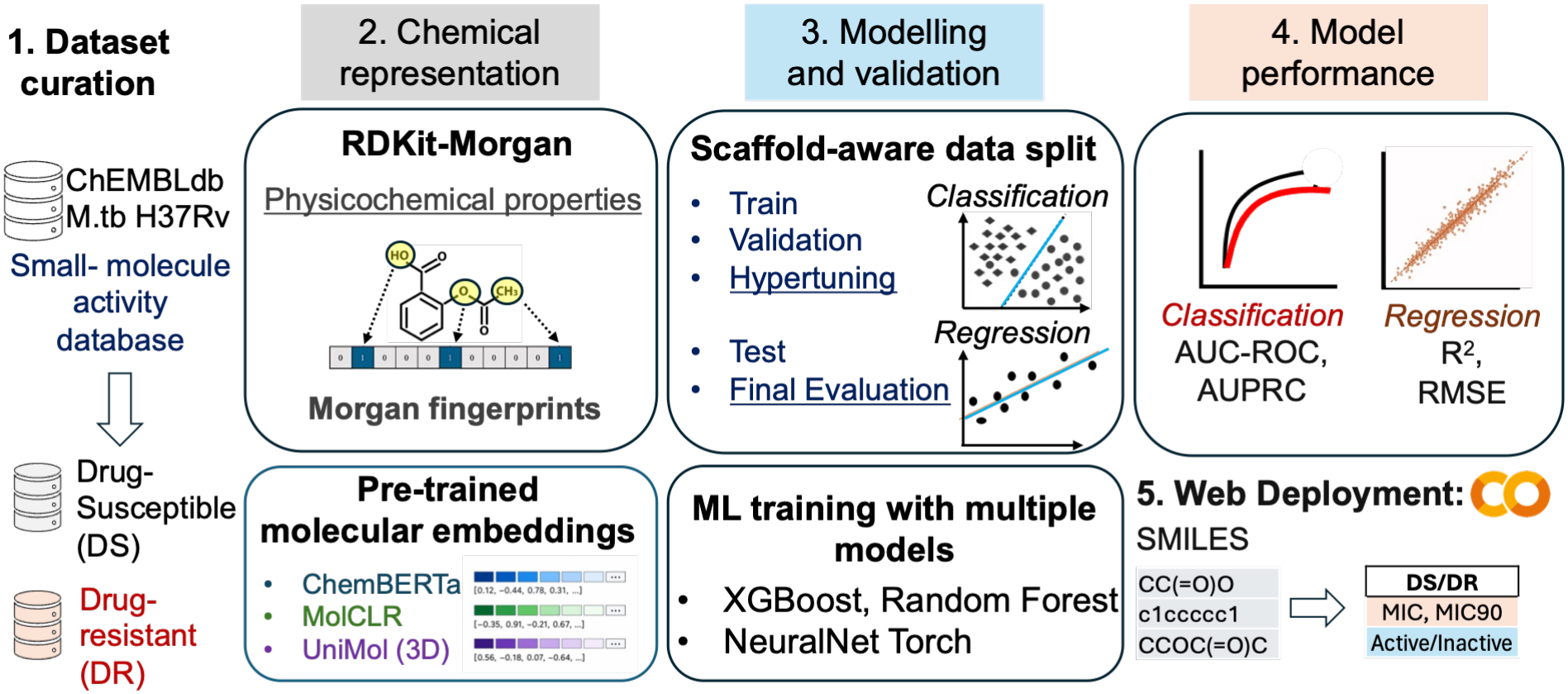
MtbAIM workflow for prediction of M.tb drug susceptibility and antimicrobial activity. Bioactivity data for M.tb H37Rv were curated from ChEMBL and processed for downstream modelling. Compounds were represented using physicochemical descriptors and Morgan fingerprints, as well as learned molecular representations from ChemBERTa, MolCLR, and UniMol. Models were trained, hyperparameter-tuned, validated, and evaluated using held-out test data. Classification performance was assessed using AUROC and AUPRC, whereas regression performance was evaluated using R^2^ and RMSE. The resulting models were integrated into a web-based prediction (MtbAIM) framework accepting SMILES as input to predict drug-susceptible/drug-resistant (DS/DR) activity, quantitative MIC/MIC90 endpoints, and active/inactive classifications.

## Methods

### Initial data retrieval and curation

Whole cell anti-mycobacterial bioactivity records were obtained and curated from ChEMBL (version 37)^6^. Records were filtered specifically for M.tb H37Rv, excluding H37Ra and non-M.tb organisms. DS and DR records were maintained as separate datasets for downstream analysis.

### Replicate aggregation and regression targets

Replicate measurements were aggregated only within the same canonical structure, biological context, and endpoint (no pooling of MIC, MIC90, and MIC99 endpoints). The median activity was used as the compound-endpoint summary to reduce sensitivity to individual extreme measurements. Quantitative activities were transformed to pMIC as −log10(MIC in mol L^−1^). After endpoint-specific filtering and structure quality control, the regression datasets contained 21,683 DS MIC, 3,754 DS MIC90, and 1,846 DS MIC99 compounds, and 3,220 DR MIC, 561 DR MIC90, and 180 DR MIC99 compounds (Supplementary Figure 1). All of these six datasets were modeled independently.

### Binary activity definition

For binary classification, a deliberate activity boundary was used to reduce ambiguity around intermediate-potency compounds (Supplementary Figure 1). Records with aggregated activity <5,000 nM were assigned to the active class and records >15,000 nM to the non-active class. After final structure and label quality control, the classification datasets contained 25,645 DS compounds (annotated as 12,591 active and 13,054 non-active) and 3749 DR compounds (annotated as 2,215 active and 1,534 non-active).

### Molecular representations

For a rigorous comparative analysis, four different molecular representations were evaluated. The classical representation combined 217 RDKit physicochemical/topological descriptors with a 2048-bit Morgan fingerprint (radius 2, chirality enabled), giving 2265 input features per compound^13, 14^. Learned representations were generated from three pretrained molecular models. ChemBERTa used the DeepChem/ChemBERTa-77M-MLM checkpoint and mean pooling over SMILES tokens to obtain a 384-dimensional embedding^10^. MolCLR^11^ generated 512-dimensional hidden-state embeddings from a pretrained GIN graph encoder, and UniMol generated 512-dimensional molecular representations with UniMol^12^. All embeddings were generated independently of the compound activity labels.

### Scaffold-aware cross-validation

First, generalization was evaluated using five-fold Bemis-Murcko scaffold-aware cross-validation^15^. For the binary task, 8,601 unique DS scaffolds and 1,470 unique DR scaffolds were assigned to frozen master folds, which were reused across RDKit/Morgan, ChemBERTa, Uni-Mol, and MolCLR analyses. No scaffold was permitted to occur in both the training and held-out partition of a fold. Regression used the same principle but generated to an endpoint-specific scaffold (MIC, MIC90, and MIC99). Within each dataset-endpoint combination, these folds were frozen and reused across all representations and model families. The resulting regression sets contained 7001, 1596, and 631 DS scaffolds for MIC, MIC90, and MIC99, respectively, and 1221, 240, and 98 DR scaffolds. This design prevents close scaffold analogues from being distributed across training and test data and provides a more stringent estimate of performance on chemically distinct compounds.

### Model development and hyperparameter optimization

Classical classification models were first screened within each representation, after which the leading candidates were optimized separately in the five outer scaffold folds. Candidate algorithms included ExtraTrees, Random Forest, histogram gradient boosting, XGBoost, LightGBM, and CatBoost, with the subset taken forward determined by the representation and dataset. Hyperparameters and the classification decision threshold were selected using only the training portion of each outer fold where Optuna was used for the focused tuning stage^16^. Balanced accuracy was the principal threshold-dependent selection criterion, with Matthews correlation coefficient (MCC), AUROC, sensitivity, specificity, precision, and AUPRC retained for complementary assessment. In parallel, multilayer perceptrons were trained on ChemBERTa, UniMol, and MolCLR embeddings. Four predefined network architectures were evaluated within each outer fold, with early stopping used during training (maximum 100 epochs and patience 10 for classification).

### Endpoint-specific regression modeling

Regression was performed separately for each of the six DS and DR endpoint datasets. Initial benchmarking evaluated Ridge and Elastic Net regression together with Random Forest, ExtraTrees, histogram gradient boosting, XGBoost, LightGBM, and CatBoost across all four molecular representations. Hyperparameter optimization was then applied to the strongest representation-model combinations using the frozen endpoint-specific scaffold folds. Further, neural regression models were evaluated on ChemBERTa, UniMol, and MolCLR embeddings using four multilayer-perceptron architectures, a maximum of 150 epochs, and early stopping with patience 15. Model selection was based on held-out scaffold performance, with RMSE as the primary error metric and MAE and R^2^ reported as complementary measures.

### Performance estimation and statistical comparison

Classification performance was summarized as the mean and standard deviation across the five outer scaffold folds and from pooled out-of-fold predictions. AUROC and AUPRC quantified ranking performance, while balanced accuracy and MCC were emphasized for threshold-dependent evaluation^17, 18^. Precision-recall performance was interpreted relative to the active-class prevalence. Regression performance was evaluated from pooled out-of-fold RMSE, MAE, and R^2^, together with fold-level variability and bootstrap confidence intervals for final models. Classical and neural regression finalists were additionally compared using paired out-of-fold errors, with false-discovery-rate correction applied across the endpoint comparisons. All representation and model comparisons within a dataset used the same frozen scaffold partitions, ensuring that performance differences reflected the molecular representation/model combination rather than different train-test splits.

## Results

In this work, we have established a rigorously curated and chemically diverse dataset to systematically evaluate ML approaches for antitubercular activity prediction. Using scaffold-aware cross-validation, we compared classical molecular descriptors and fingerprints with pretrained molecular representations across both DS and DR M.tb. We then evaluated these representations across binary classification and endpoint-specific regression tasks to determine their ability to generalize to structurally distinct compounds.

Classification performance differed substantially across molecular representations (Figure 2). In the DS dataset, the strongest classical model was ExtraTrees trained on RDKit/Morgan features, with a mean balanced accuracy of 0.782±0.010, AUROC of 0.864±0.007, and AUPRC of 0.870±0.008 across the five scaffold folds. Alternatively, the best neural model using ChemBERTa embeddings reached a balanced accuracy of 0.738±0.009, AUROC of 0.813±0.013, and AUPRC of 0.822±0.012. The DR dataset showed a similar pattern, where RDKit/Morgan with histogram gradient boosting achieved highest balanced accuracy (0.762±0.023; AUROC 0.836±0.027; AUPRC 0.880±0.026), while the ChemBERTa multilayer perceptron was the strongest neural comparator (balanced accuracy 0.749±0.017; AUROC 0.830±0.026; AUPRC 0.867±0.027). Corresponding, threshold-dependent balanced accuracy, MCC, sensitivity, and specificity are provided in Supplementary Figure 2.

**Figure 2.**
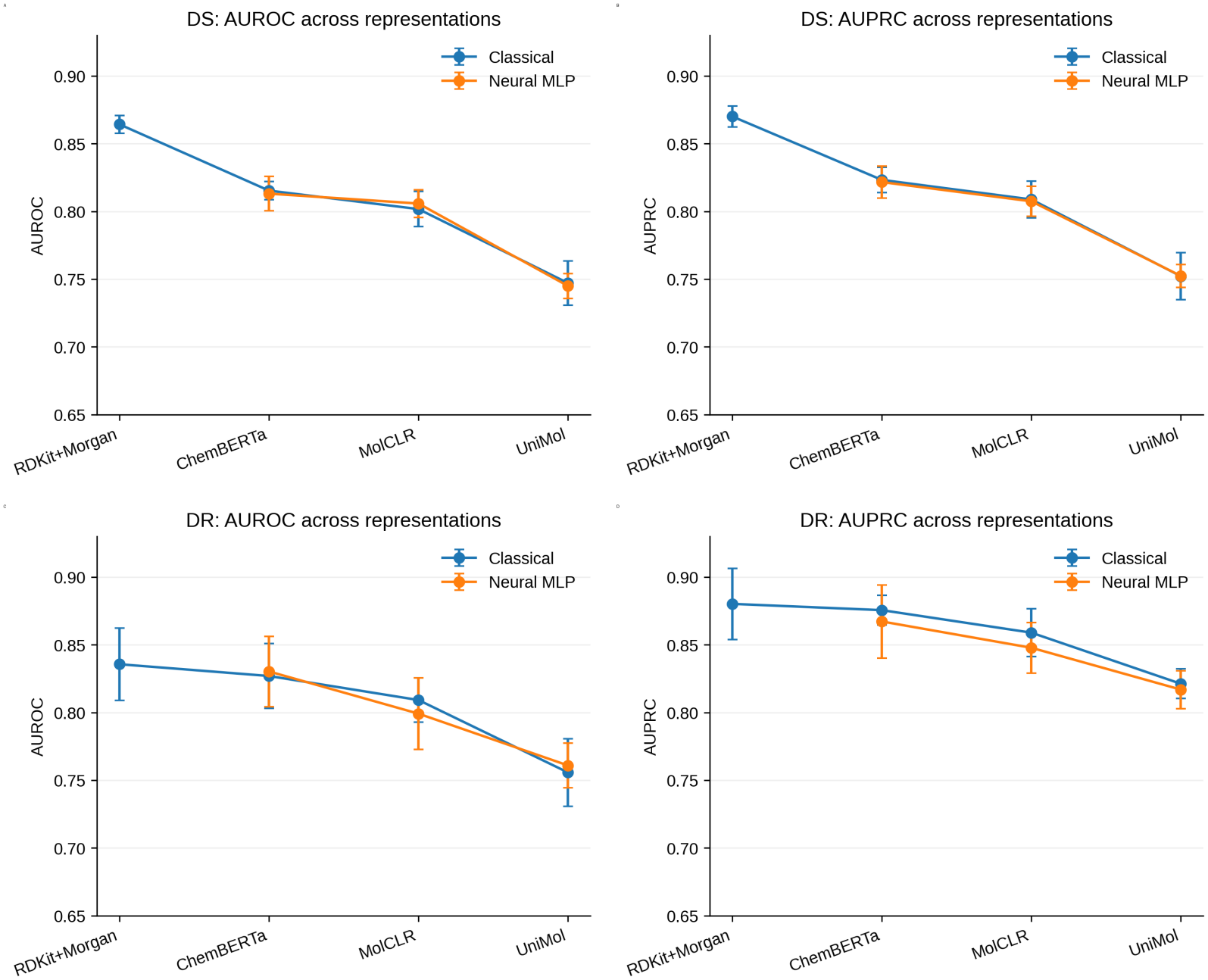
Scaffold-aware classification performance across molecular representations. Cross-validated AUROC and AUPRC for drug-susceptible (DS) and drug-resistant (DR) M.tb activity classification using RDKit/Morgan fingerprints, ChemBERTa, MolCLR, and UniMol representations. Classical values correspond to the best tuned classifier for each representation; neural values correspond to multilayer perceptron models trained on learned molecular embeddings. Points show mean performance across five outer scaffold folds and error bars show standard deviation. RDKit/Morgan was evaluated only with classical models.

Pooled out-of-fold ROC and precision-recall curves also corroborated with previous observations (Figure 3), where class discrimination was retained after combining predictions from all five held-out scaffold folds. This is particularly important for the DR dataset, where the number of compounds is much smaller and fold-to-fold variation is consequently greater. The precision-recall curves were interpreted against the active-class prevalence rather than against zero, providing a more useful reference for the imbalanced setting^19^.

**Figure 3.**
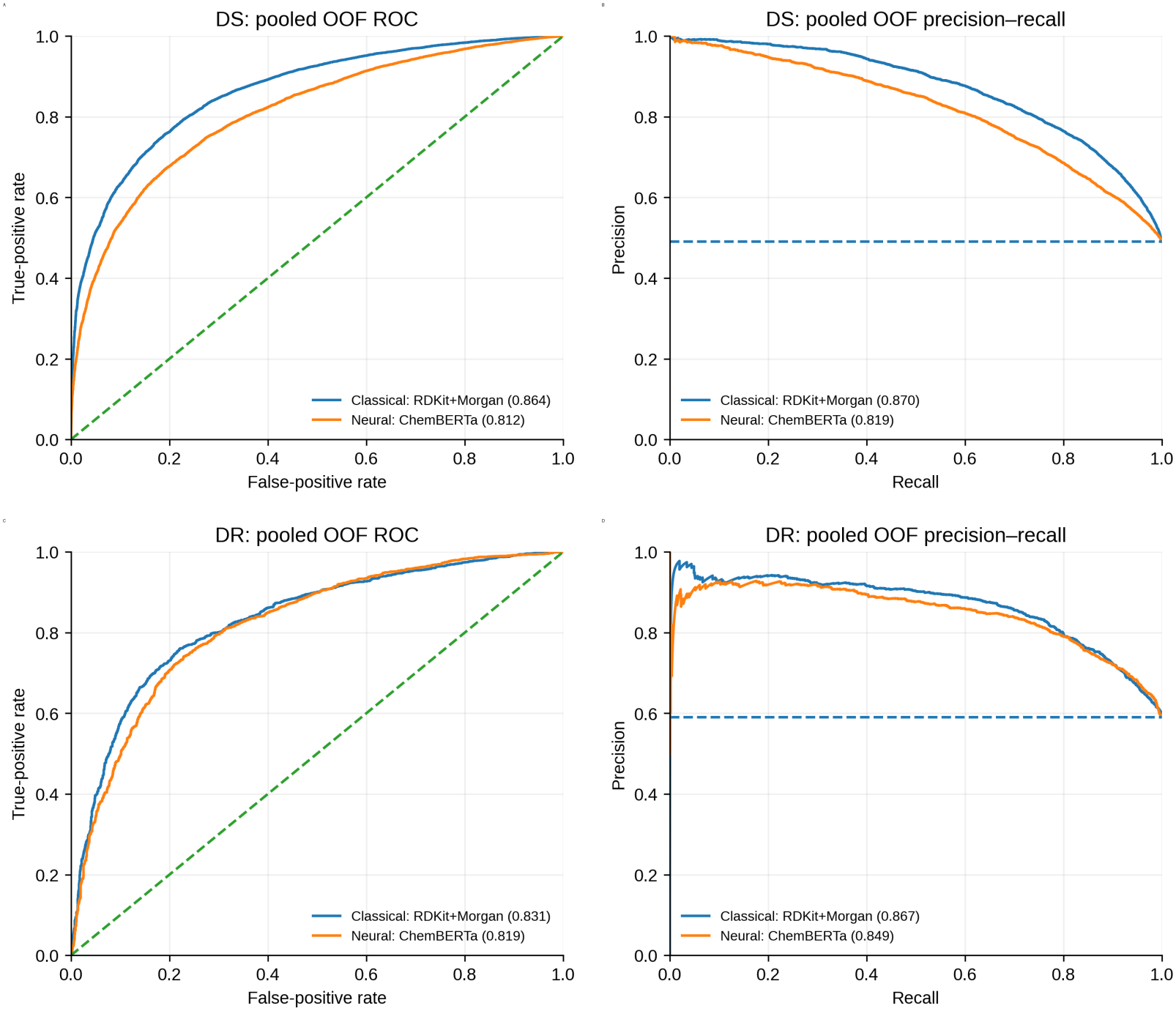
Out-of-fold discrimination of active and non-active compounds. Pooled out-of-fold receiver-operating characteristic (ROC) and precision-recall curves for the best classical and neural classification models in DS and DR datasets. Classical curves correspond to the highest-balanced-accuracy tuned classifier in each dataset (RDKit/Morgan with ExtraTrees for DS and histogram gradient boosting for DR); neural curves correspond to ChemBERTa-MLP models. Diagonal ROC reference lines indicate random discrimination; horizontal precision-recall references indicate the active-class prevalence in each dataset. Values in legends are pooled OOF AUROC or AUPRC.

In our second analysis of anti-mycobacterial structure-activity data, endpoint-specific regression produced useful quantitative separation across all six DS/DR datasets (Figure 4). For DS dataset, the best models gave pooled out-of-fold RMSE/R^2^ values of 0.714/0.490 for MIC, 0.682/0.601 for MIC90, and 0.670/0.541 for MIC99. For DR dataset, the RMSE/R^2^ for corresponding MIC, MIC90 and MIC99 values were 0.729/0.546, 0.784/0.613, and 0.604/0.790. The best performing models for DS datasets were ExtraTrees for MIC, LightGBM for MIC90, and XGBoost for MIC99, and ExtraTrees for all three DR endpoints. However, for every endpoint-dataset combination, classical RDKit/Morgan features performed slightly better than molecular embeddings. The observed-versus-predicted distributions are shown in Figure 5, while representation-level benchmarks and fold-to-fold RMSE variability are provided in Supplementary Figures 3 and 4.

**Figure 4.**
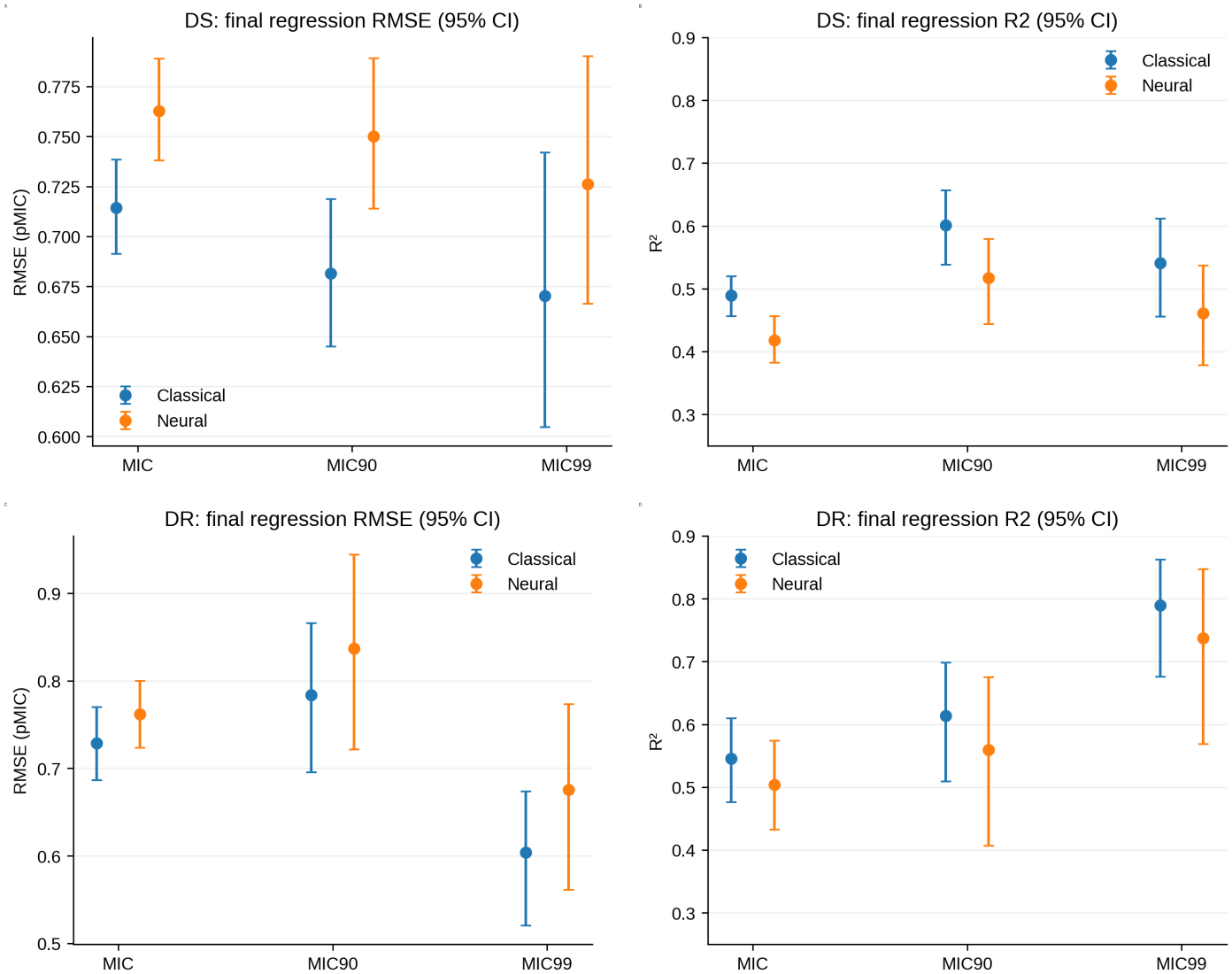
Final endpoint-specific pMIC regression performance under scaffold-held-out validation. RMSE and R^2^ of the best classical and best neural regression finalists for MIC, MIC90, and MIC99 in DS and DR datasets. Points denote pooled out-of-fold estimates and error bars denote 95% scaffold-bootstrap confidence intervals from the final comparison stage. Across all six endpoint-dataset combinations, the best overall model used RDKit/Morgan fingerprints with a tree-based regressor.

**Figure 5.**
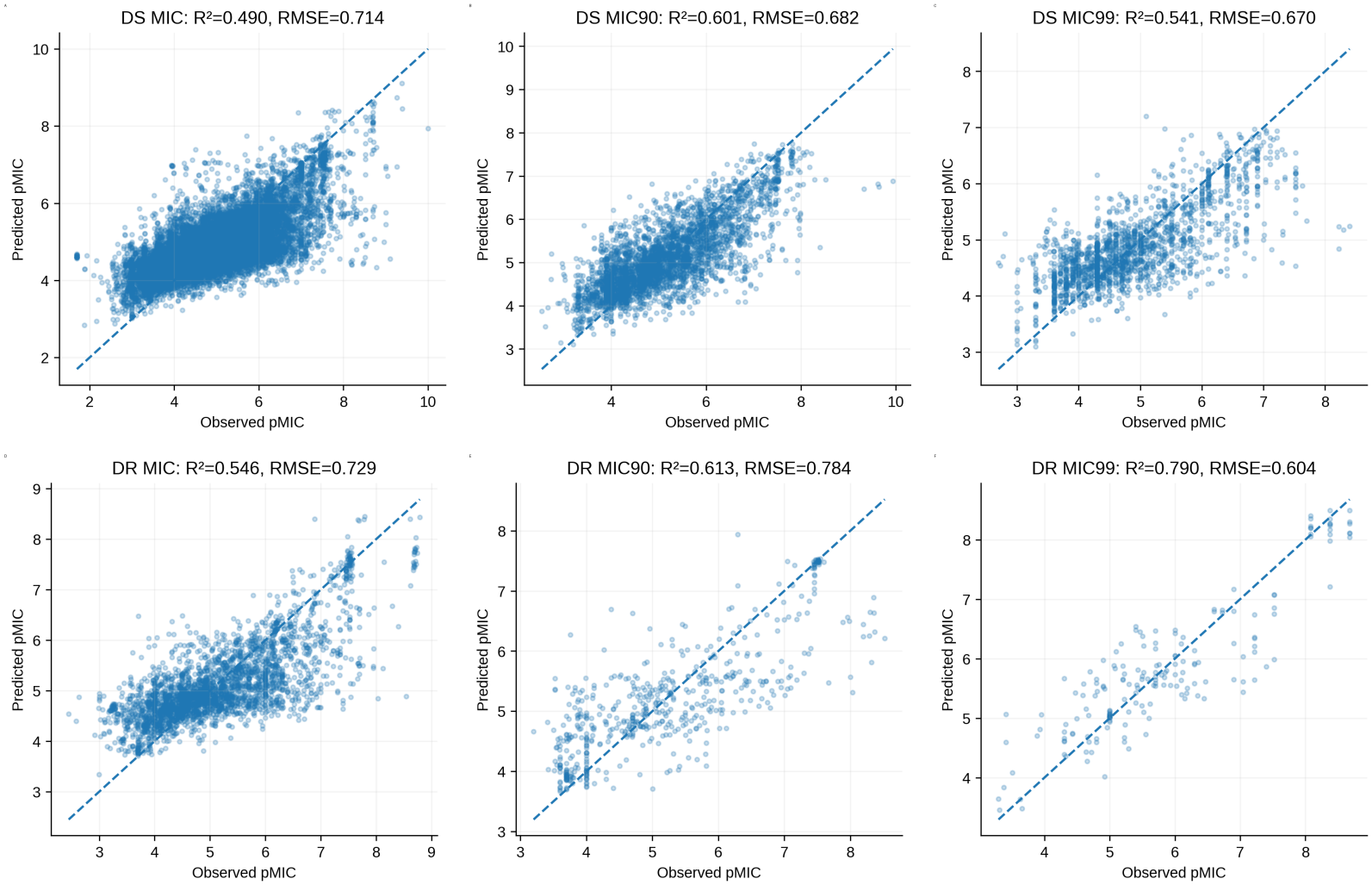
Observed-versus-predicted pMIC values for final best-overall regression models. Parity plots of pooled out-of-fold predictions from the final best-overall model for DS and DR MIC, MIC90, and MIC99 datasets. Dashed lines indicate the identity relationship. Panel titles report pooled OOF R^2^ and RMSE. Points represent scaffold-held-out compounds; underlying compound-level predictions are provided in the associated raw-data CSV.

The regression comparison also indicated that stronger model complexity did not automatically translate into better scaffold-held-out performance. In DS, the best neural model had RMSE values higher than the best classical model by 0.048, 0.069, and 0.056 pMIC for MIC, MIC90, and MIC99, respectively; all three differences remained significant after false-discovery-rate correction. In DR, the RMSE difference was smaller for MIC (+0.033 pMIC) and was not statistically resolved for MIC90 or MIC99. Prediction accuracy expressed as 2-, 4-, and 8-fold concentration error is summarized in Supplementary Figure 5, and the paired classical-versus-neural comparisons are shown in Supplementary Figure 6.

### Finalized prediction models and MtbAIM platform

Based on the comparative evaluation, eight models were selected for deployment in MtbAIM. ExtraTrees and histogram gradient boosting were retained for DS and DR binary classification, respectively. For regression, ExtraTrees was selected for DS MIC and all three DR endpoints, while LightGBM and XGBoost were selected for DS MIC90 and MIC99, respectively. All finalized models used the RDKit/Morgan molecular representation.

These models were integrated into a unified prediction workflow accessible through Google Colab (https://github.com/jasdeep002/MtbAIM). The platform accepts individual SMILES or compound lists in TXT/CSV format and supports batch predictions of up to 100 compounds per submission. Users can obtain DS- and DR-specific activity classifications, predicted potency values for individual MIC endpoints, and associated prediction intervals in downloadable CSV files. This implementation allows the selected models to be applied to new chemical structures without requiring users to perform molecular feature generation or model training independently.

## Discussion

Tuberculosis drug discovery remains challenging due to continuing global disease burden, and drug resistance further complicating the identification and prioritization of effective compounds. The 2025 WHO Global Tuberculosis Report highlights drug-resistant TB as a persistent public-health threat and emphasizes the continuing need for research, innovation, drug-susceptibility testing, and development of new therapeutic tools^1^. Computational approaches have increasingly been used to prioritize antitubercular small molecules, but most QSAR (Quantitative Structure Activity Relationship) and machine-learning studies have focused predominantly on activity against the drug-susceptible M.tb or using individual chemical series^20^. Some QSAR studies have extended this by experimentally evaluating prioritized compounds against resistant strains, demonstrating the value of resistance-aware screening. For example, QSAR-guided chalcone design identified compounds active against H37Rv as well as rifampicin- and isoniazid-resistant strains, while another ANN/QSAR study identified compounds active against both H37Rv and INH/RIF-resistant *M. tuberculosis*^21^. More recently, an ML/deep-learning virtual-screening workflow screened 11,576 compounds and experimentally identified candidates active against both H37Rv and multidrug-resistant isolates; however, resistant-isolate activity was used primarily for downstream validation rather than being modeled as a distinct predictive platform^22^. Several computational platforms have been developed to support small-molecule prioritization against M.tb. Tools such as ChemTB, mycoCSM, and Assay Central-based M.tb models demonstrate that machine learning can successfully identify or rank compounds with predicted antitubercular activity^20, 23, 24^, while platforms such as the Ersilia Model Hub have improved accessibility of deployable molecular models^25^. However, these approaches generally emphasize overall M.tb activity, H37Rv-focused prediction, or single-task screening and do not explicitly treat drug-susceptible and drug-resistant isolates as separate predictive domains. In contrast, the present framework (MtbAIM) was designed to preserve this distinction and to combine endpoint-specific classification with quantitative regression, allowing prioritization not only on the basis of predicted activity but also on whether activity is expected to be retained across DS and DR M.tb. The DR models also address a different problem from genomic resistance prediction. Sequence-based diagnostic models estimate whether an isolate is resistant to an existing drug^19^. Here, the input is the chemical structure, and the question is whether that structure is likely to inhibit an isolate already described as drug resistant. The two approaches are therefore complementary: one supports resistance diagnosis, whereas the other is intended to help prioritize new or repurposed chemistry for experimental testing against resistant M.tb. A second important finding in our study was that the learned molecular representations were informative but did not consistently outperform the classical RDKit/Morgan representation. ChemBERTa was the strongest neural comparator for both DS and DR classification and for most regression tasks, yet the final best-overall models were still based on RDKit/Morgan features. The paired regression analysis made this difference clearest in the larger DS datasets, where the classical advantage remained significant for all three endpoints. The endpoint-specific results also highlight the role of data density. MIC provided the largest training sets in both isolate contexts, whereas MIC90 and especially MIC99 became progressively smaller. The DR MIC99 model, for example, produced a high pooled R^2^ but was evaluated on only 180 compounds. Its estimate is therefore less stable than the headline value alone might suggest. Keeping the endpoints separate makes this uncertainty visible; pooling them would increase sample size but would hide the different levels of experimental support. Collectively, the intended use of MtbAIM is compound prioritization, not replacement of microbiological testing. Public bioactivity data remain heterogeneous in assay conditions, strain annotation, censoring, and measurement density. While scaffold-aware cross-validation reduces analogue leakage, prospective temporal or independently generated experimental data will be needed to establish performance on truly unseen chemical series. Within those limits, the combined classification and regression outputs provide a practical way to rank compounds by predicted activity, isolate context, and expected potency before committing to resource-intensive M.tb assays.

## Conclusion

Separating isolate susceptibility and MIC-family endpoint produced a more specific view of antitubercular activity than a single pooled model. Across scaffold-held-out validation, RDKit/Morgan features with tree-based learners gave the strongest overall classification and regression performance, while ChemBERTa, MolCLR, and Uni-Mol captured substantial but not consistently superior signal. The resulting models provide complementary predictions for drug-susceptible and drug-resistant M.tb and for MIC, MIC90, and MIC99 potency. This structure is well suited to compound prioritization because it keeps the biological context of the experimental measurement visible in the prediction.

## Supporting information

and 3,220 DR MIC, 561 DR MIC90, and 180 DR MIC99 compounds

## Code and model availability

MtbAIM is publicly accessible through GitHub (https://github.com/jasdeep002/MtbAIM). The Google Colab implementation provides access to the finalized classification and regression models, enabling prediction of antitubercular activity and endpoint-specific potency from user-supplied molecular structures.

## Acknowledgements and Funding statement

The authors thank the Indian Institute of Technology-Delhi (India) for infrastructural support. J.S. acknowledges Young Scientist Scheme (Grant number: 12014/16/2018-HR) of Dept. of Health Research, Ministry of Health & Family Welfare, Govt of India (2018-22). K.S. acknowledges fellowship support from SERB-NPDF (Sanction No: PDF/2018/004414). Authors also acknowledge that during the preparation of this manuscript, they used AI assistance to help refine the grammar, and clarity. The content was carefully reviewed, edited, and authors take full responsibility for the final content.

