## Supplementary material for "MtbAIM: AI/ML-guided endpoint-specific prediction of small-molecule activity across drug-susceptible and drug-resistant *Mycobacterium tuberculosis*": and 3,220 DR MIC, 561 DR MIC90, and 180 DR MIC99 compounds


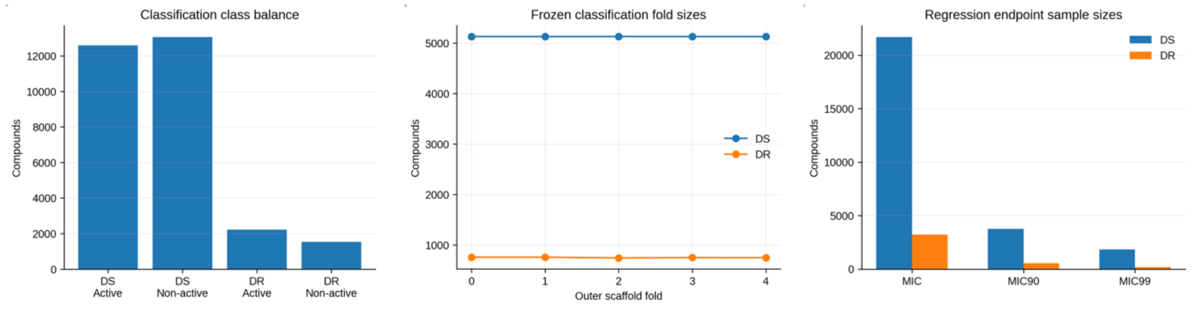


**Supplementary Figure 1. Dataset composition and scaffold-split quality control.** Class balance in the DS and DR binary classification datasets, compound counts across the five frozen classification scaffold folds, and final compound counts for MIC, MIC90, and MIC99 regression endpoints.

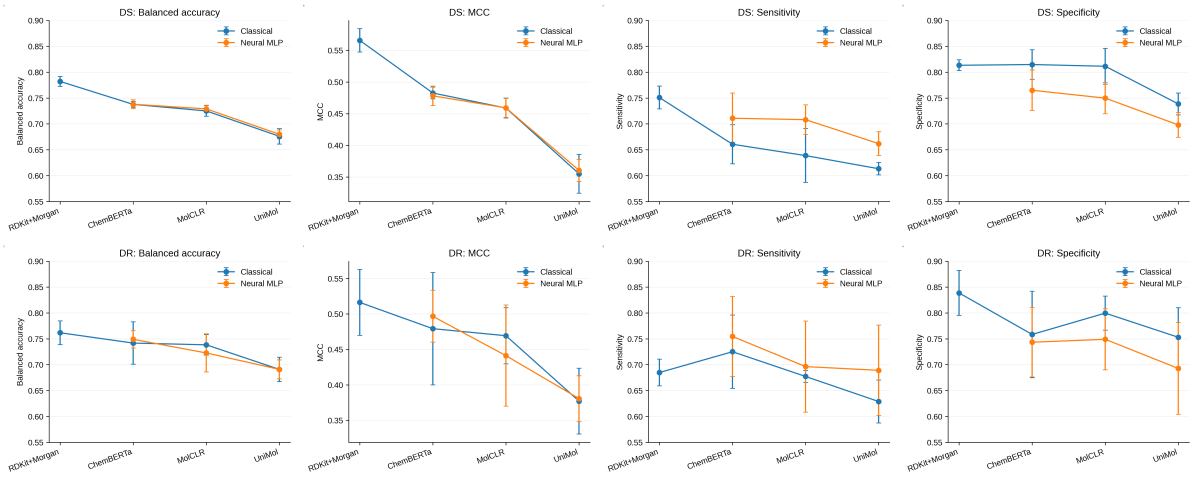


**Supplementary Figure 2. Threshold-dependent classification metrics across representations.** Balanced accuracy, Matthews correlation coefficient, sensitivity, and specificity for the best classical classifier and neural multilayer perceptron for each representation in DS and DR datasets. Points show five-fold means and error bars show the standard deviation.

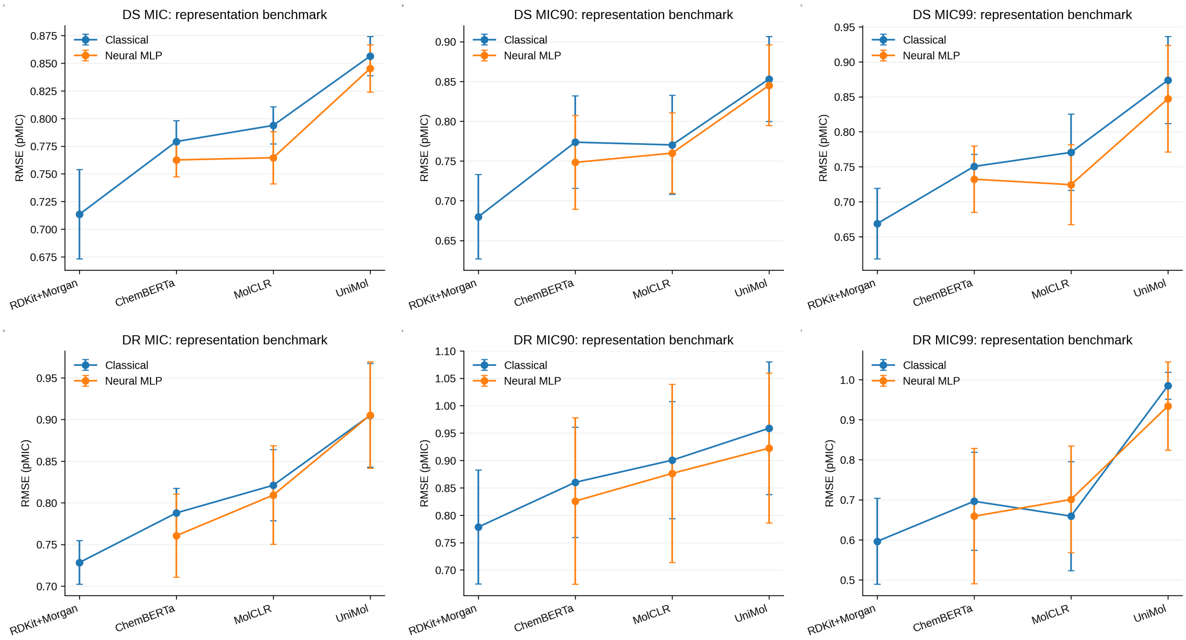


**Supplementary Figure 3. Representation-level regression benchmark.** Five-fold RMSE for the best classical regressor and neural multilayer perceptron associated with each molecular representation for all DS and DR endpoints. The comparison shows the performance of RDKit/Morgan, ChemBERTa, MolCLR, and Uni-Mol before final classical-versus-neural finalist selection.

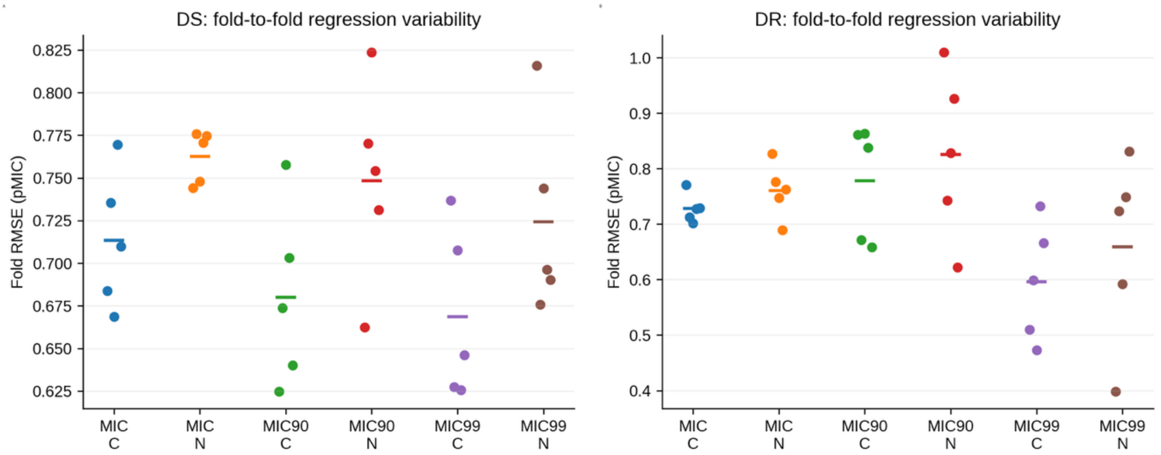


**Supplementary Figure 4. Fold-to-fold variability of final regression finalists.** Outer-fold RMSE distributions for the best classical and neural finalist at each DS and DR endpoint. Individual points show the five scaffold folds and horizontal markers show fold means.

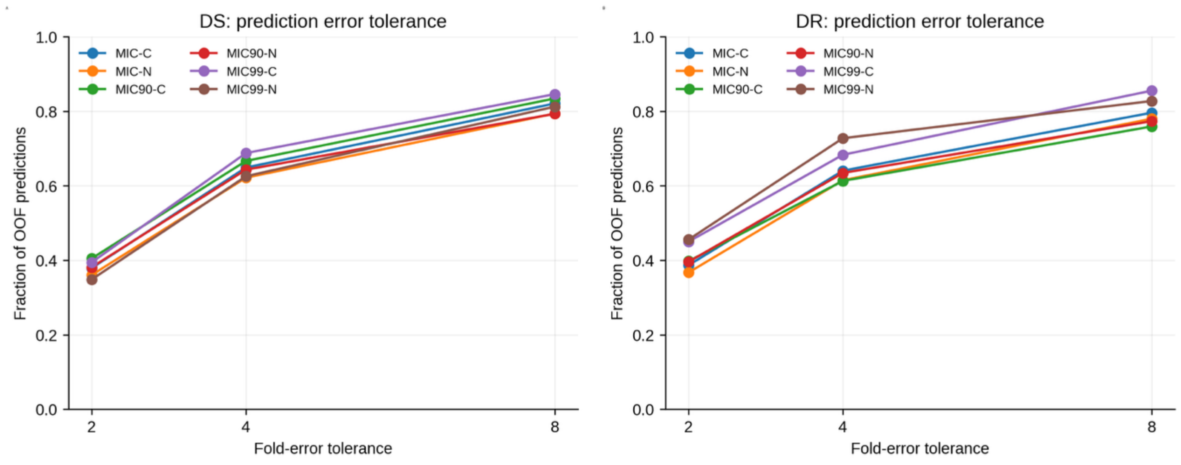


**Supplementary Figure 5. Prediction accuracy expressed as fold-error tolerance.** Fraction of pooled OOF predictions falling within 2-fold, 4-fold, and 8-fold concentration error for the best classical (-C) and neural regression (-N) finalists across MIC, MIC90, and MIC99 endpoints.

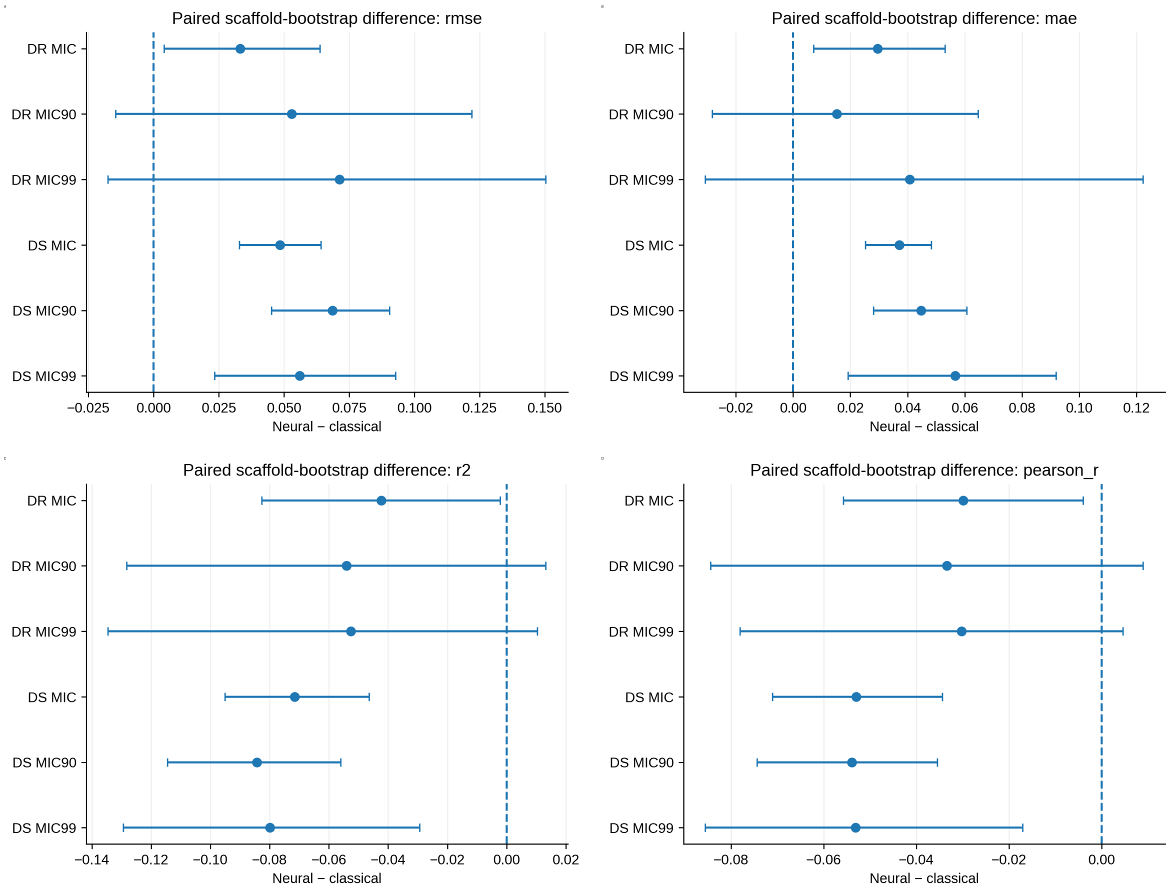


**Supplementary Figure 6. Paired classical-versus-neural regression comparisons.** Paired neural-minus-classical differences for RMSE, MAE, R2, and Pearson correlation across all dataset-endpoint combinations. Error bars show the reported paired confidence intervals; zero denotes no difference. FDR-adjusted P values and significance calls are provided in the associated raw-data table.

Table S1. **Finalized machine-learning models selected for deployment in MtbAIM.** Summary of the eight locked models for binary activity classification and endpoint-specific MIC, MIC90, and MIC99 regression against drug-susceptible (DS) and drug-resistant (DR) *Mycobacterium tuberculosis* isolates, including the selected algorithms and molecular representations.

| Prediction task | Final model | Molecular representation |
|---|---|---|
| DS classification | ExtraTreesClassifier | RDKit/Morgan |
| DR classification | HistGradientBoostingClassifier | RDKit/Morgan |
| DS MIC regression | ExtraTreesRegressor | RDKit/Morgan |
| DS MIC90 regression | LightGBMRegressor | RDKit/Morgan |
| DS MIC99 regression | XGBoostRegressor | RDKit/Morgan |
| DR MIC regression | ExtraTreesRegressor | RDKit/Morgan |
| DR MIC90 regression | ExtraTreesRegressor | RDKit/Morgan |
| DR MIC99 regression | ExtraTreesRegressor | RDKit/Morgan |
